# Model of naturally occurring refractive error (NORE) in mice

**DOI:** 10.64898/2026.07.01.735855

**Authors:** Melissa R. Bentley-Ford, Teele Palumaa, Linjiang Lou, Aparna Jonnalagadda, Morgan L. Bade, Shruti Balamurugan, Reece Mazade, Machelle T. Pardue

## Abstract

**Purpose:** Animal models of myopia typically induce monocular refractive shifts via form deprivation (FD) or lens-induced myopia (LIM), modeling susceptibility to myopia, but with potentially limited applicability to childhood myopia. Here we describe a novel, genetically diverse mouse model of naturally occurring refractive error (NORE) with three distinct refractive phenotypes: hyperopic, myopic, and intermediate.

**Methods:** C57BL/6J mice were mated to 129S2/SvPasCrl mice to create F1 or F2 offspring. Refractive errors in male and female F1 (N=21) and F2 (N=101) mice were assessed on postnatal days (P) 28 and 42 using photorefractometry. In a subset of mice (N=30–40), corneal radius of curvature, axial ocular dimensions, retinal and visual function were assessed.

**Results:** F2 mice were classified as NORE with either hyperopic (RE≥0 diopters (D) at P28 and P42), myopic (RE<0D at P28 and P42) or intermediate (RE<0D at P28 and RE≥0D at P42) refractions based on individual trajectories. All ocular parameters changed with age, with significantly slower growth in axial length and vitreous chamber depth in the intermediate versus myopic mice (p<0.05). Lens thickness was smaller in the myopic group at P28. Differences in refraction were not attributed to variances in retinal function or dopamine signaling.

**Conclusions:** NORE mice represent a novel, genetically diverse wild-type mouse model that, unlike traditional models, does not require interventions such as FD or LIM to induce myopia. NORE mice provide a valuable tool for future investigations of genetic and environmental mechanisms and targeted therapeutic strategies for refractive errors.

## Introduction

Uncorrected refractive errors are the leading cause of visual impairment worldwide, with myopia being the most common form. The prevalence of myopia has increased at a staggering rate and continues to rise^1–3^. At its current trajectory, 40-50% of the global population is predicted to be affected by 2050^3,4^. This trend is concerning, as the risk of developing vision-threatening disorders such as retinal detachment, macular degeneration, cataracts, and glaucoma later in life increases with the severity of myopia^5^. Based on this alarming forecast, there is an urgent need for tools to better understand the mechanisms driving myopia development.

A combination of genome-wide association studies (GWAS) and twin studies has provided substantial support for the genetic contribution to myopia^6–17^. GWAS have identified more than 900 single-nucleotide polymorphisms (SNPs) associated with refractive error and over 800 SNPs associated with myopia, with one meta-analysis of European ancestry estimating that the associated GWAS variants can explain 18.4% of heritability^16^. Comparatively, twin studies consistently report a much higher contribution of genetic factors, ranging from 70-90%^14^. For a comprehensive review, see *Myopia: Causes, Prevention, and Treatment of an Increasingly Common Disease* (National Academies of Sciences, Engineering, and Medicine)^18^. Myopia frequently runs in families, with myopic parents being more likely to have children with myopia^19–23^. While this is likely due to a combination of genetic, behavioral, and environmental factors, it supports the idea that myopia results from complex interactions among these factors. Understanding how changes in a visual environment are detected by the retina and translated into chemical signaling that propagates from the retina through the posterior structures of the eye, ultimately culminating in changes to the scleral extracellular matrix and its biomechanical properties, is critical for developing safe and effective myopia management and prevention strategies.

Animal models, such as macaques, chickens, tree shrews, guinea pigs, marmosets, zebrafish, and mice, have provided invaluable mechanistic insights into myopia^24^ ^25^. Traditional animal models of refractive error use monocular form deprivation or lens defocus to induce form-deprivation myopia (FDM), lens-induced myopia (LIM), or lens-induced hyperopia (LIH)^24^. In FDM, a diffuser or a frosted lens is placed in front of one eye, reducing image clarity and driving axial elongation^26,27^. By contrast, lens defocus uses a negatively or positively powered lens to create defocus, causing light to focus behind or in front of the retina and promoting subsequent axial elongation or slowing of axial growth, respectively^24,28^ ^29^. In these models, the treated eye develops a refractive error relative to the contralateral eye over days or weeks.

While each model has its strengths and weaknesses, mice offer significant advantages given the availability of genetic tools and resources^30^. In myopia studies, it is standard practice to use C57BL/6J wild-type mice, which naturally develop a hyperopic refractive error with age^31^. Mice respond to FD and lens defocus, like other animal models, although changes in axial length can be difficult to measure due to the small size of the mouse eye (1D optical power change = ∼6 microns of axial change)^30,32^. In addition to induced myopia models, transgenic mouse models have highlighted several genes and pathways that are risk factors for refractive errors or myopia susceptibility (for a complete list of genes, see^30^). For instance, *Nyx*-mutant mice have more hyperopic refractive errors during normal development but are more susceptible to form deprivation myopia than wild-type mice^33^. Additionally, different inbred strains of mice have distinct refractive development curves. For example, substrains of 129 mice have myopic refractive errors in the first few weeks following birth, whereas the C57BL/6 strain is hyperopic^34,35^.

In this study, we leverage the natural variation between 129S2/SvPasCrl (129) and C57BL/6J (C57) strains to generate an F2 cross that produces a novel naturally occurring refractive error (NORE) mouse model. The NORE model captures the genetic diversity and variability in refractive development necessary to study the complex mechanisms underlying myopia.

## Methods

### Animals

Mice were housed in a 12:12-hour light: dark cycle in standard animal facility lighting (30-240 lux in a standard cage) at Emory University School of Medicine. All procedures were approved by the Emory University Institutional Animal Care and Use Committee and adhered to the ARVO Statement for the Use of Animals in Ophthalmic and Vision Research. C57BL/6J mice (Jackson Labs, Bar Harbor, ME, USA) and 129S2/SvPasCrl (Charles River, Wilmington, MA, USA) mice were mated together (F0) to generate an isogenic F1 population of mice, which were subsequently mated together to generate the F2 mice used for all experiments. It was confirmed that mating pairs were not carriers of retinal degeneration point mutations such as *Rd1, Rd8,* or *Rd10*.

### Experimental design

The initial timepoint for refractometry, keratometry, and OCT biometry measurements was performed on male and female mice at P29 (+/- 1 day). Visual acuity, ERG response, and contrast sensitivity were assessed at P40, P41, and P42 days, respectively. Additional refractometry, keratometry, and OCT biometry measurements were performed again 14 days after the initial measurements (P42-44). Experimental F2 mice consisted of at least 20 litters from multiple F1 mating pairs, and F1 mating pairs were generated from multiple F0 mating pairs.

### Photorefractometry and keratometry measurements

After dilating the eyes with 1% tropicamide and confirming dilation, mice were anesthetized with a ketamine/xylazine cocktail (100mg/10/kg) based on their body weight. Refractive error (RE) was measured using an automated eccentric infrared photorefractor as described previously^31,33^. After refractive measurements, the anterior corneal radius of curvature was measured using an automated infrared keratometer^31,36^.

### Ocular biometrics

Ocular axial dimensions were measured using spectral domain-optical coherence tomography (SD-OCT) (Envisu R4310, Leica Microsystems, Wetzlar, Germany). Three whole-eye images and one retinal scan were taken of each eye at each time point. For whole-eye images, one radial scan (0.4 mm, 1000 A-scans/B-scan, 20 B-scans, 1 frame/B-scan) and two linear scans (0.4 mm, 1000 A-scans/B-scan, 10 B-scans, 2 frames/B-scan) were captured. Scans were centered at 10 degrees superior to the optic nerve head. For retinal scans, the scan protocol included a 1-mm diameter annular scan (1000 A-scan/B-scan, 2 B-scans, 48 frames/B-scan) centered at the optic nerve head. Optical distances were converted to geometric distances using an average refractive index of 1.43^32,37^.

Whole-eye images were analyzed with a custom Python program. Tissue interfaces were manually delineated at the corneal light reflex to determine axial length, retinal thickness, vitreous chamber depth, lens thickness, anterior chamber depth, and corneal thickness. Axial length was defined as the distance from the anterior cornea to the retinal pigmented epithelium. Retinal scans were analyzed with a custom MATLAB program. The inner limiting membrane, Bruch’s membrane, the choroid/sclera interface, and the posterior border of the sclera were manually segmented. Thickness values were computed at 0.01 mm intervals across the entire scan length and averaged to calculate retinal, choroidal, and scleral thickness.

### Electroretinogram (ERG)

Retinal function was measured via full-field response to flash stimuli (UTAS BigShot; LKC Technologies, Gaithersburg, MD; differentially amplified at 1-1500 Hz with a flash duration of 5ms and a sampling rate of 2000 Hz) of increasing luminance (-3 to 2.5 log cd s/m^2) in animals that were dark-adapted for at least 4 hours and prepared under dim red light conditions. The pupils were dilated with 1% tropicamide, and animals were anesthetized with a ketamine/xylazine cocktail (100 mg/10 kg ketamine/ 5-10 mg/kg xylazine via intraperitoneal injection). Recording electrodes consisted of gold loop electrodes placed on the eye, referenced to a needle electrode inserted subcutaneously below the eye, and grounded to a needle electrode in the tail. ERGs were analyzed as previously reported^38,39^. The a-wave, indicating photoreceptor function,^40,41^ was measured from the baseline to the trough; the b-wave, indicating ON bipolar cell activation^42^, was measured from either the baseline or the trough of the a-wave (when present) to the waveform peak.

### Optomotor response (OMR)

A virtual OMR system (OptoMotry system, Cerebral Mechanics, Lethbridge, AB, Canada) was used to test spatial-frequency thresholds and contrast sensitivity^43,44^. Briefly, mice were placed on a five-cm-diameter pedestal in the middle of an arena surrounded by four computer-controlled 17-inch LCD screens and allowed to move freely. To determine spatial frequency thresholds, a spatial-frequency staircase was used (100% contrast, measured over 10 reversals), and for contrast sensitivity, a contrast staircase was used (0.064 cycles/degree (d), 25% contrast step size, measured over 10 reversals). These stimuli consisted of vertical sine-wave gratings that rotated at a constant speed of 12 d/s in both clockwise and counterclockwise directions. Animal movements were monitored via an overhead camera with the image projected onto a computer screen. An observer manually adjusted the designated center of focus based on the animal’s head location in real time. This ensured that the stimulus perceived by the animal was adjusted based on head and body movements^43,45^. The head turn direction was also determined by manual detection. During experimentation, the observer was blinded to the mice’s phenotype (i.e., hyperopic, myopic, or intermediate).

### Analysis

Statistical analysis was performed using GraphPad Prism 10 (San Diego, CA, USA). Unless otherwise noted, data are presented as mean ± SD, and statistical significance is denoted as *p<0.05, **p<0.01,***p<0.001, and****p<0.0001. For all datasets, ROUT outlier tests were performed. (Q=5%, GraphPad Prism 8), and any outliers were excluded. The animal order was randomly selected during experiments for the initial photorefraction, keratometry, and biometry measurements performed at P28. Subsequently, experiments were performed in the same animal order. ERG and OMR experiments were performed in a blinded manner, as final phenotype designations were completed following photorefractometry at P42. Analyses of OCT and ERG markings and HPLC samples were performed by an individual who was blinded to their respective phenotype. Figures were generated using Adobe Illustrator.

## Results

### F2 outcrossed mice resulted in a mix of refractive error (RE) phenotypes

Using the breeding strategy depicted in **Figure 1A**, the resulting F2 progeny resulted in three distinct groups: either a hyperopic (RE≥0 Diopters (D) at P28 and RE≥0 D at P42), myopic (RE<0 D at P28 and RE<0 D at P42), or intermediate (RE<0 D at P28 and RE≥0 D at P42) refractive error phenotype. At the P28 timepoint, RE significantly differed between hyperopic mice compared to myopic (+3.4±2.8D vs. -7.6±3.8D, P <0.0001, 2-way ANOVA) and intermediate mice (-5.9±2.9D, P <0.0001, 2-way ANOVA). By the P42 time point, the intermediate group rapidly progressed to hyperopic refractive errors (+4.6±2.3D). Still, it remained significantly different from both hyperopic mice (+8.7±3.6D, P <0.0001, 2-way ANOVA) and myopic mice (-4.5±2.3D, P <0.0001, 2-way ANOVA) (**Figure 1B**). Despite all three groups generally progressing towards a more positive refractive error, the change in refractive error across age in intermediate mice (+10.5±3.2D) was significantly greater than that of myopic (+3.1±4.1D, P <0.0001, Kruskal-Wallis test) and hyperopic mice (+5.8±2.9D, P=0.0006, Kruskal-Wallis test) (**Figure 1C**). Of the 155 F2 mice evaluated, 65.0% were included as experimental animals (**Figure 1D**). Anisometropia (RE differences >2D between the left and right eyes) was an exclusion criterion for 13% of animals. Additional criteria that prevented accurate measurement of RE at either P28 or P42 (i.e., cataracts, corneal defects, and small mouse size (weight less than 12.5g at P28)) affected 22% of animals and were also used as exclusionary criteria. Independent of exclusionary criteria, 51.0% of F2 progeny were female, and 49.0% were male. Overall, the breakdown of experimental F2 progeny by phenotype included 27.7% hyperopic, 30.7% myopic, and 41.6% intermediate mice, with males and females comprising 45% and 55%, overall (**Figure 1E**). Despite the ratios of male and female mice being close to the expected ratio of 50% (for both overall number of F2 mice and experimental mice), there were differences in the ratio of males and females represented within each phenotypic group: hyperopic mice: 40% males, 60% females; myopic mice: 30% males, 70% females; and intermediate mice: 60% male, 40% female (**Figure 1E)**. Ultimately, there were no significant differences in refractive development curves or progression rates across phenotypes by sex (**Supplemental Figure 1)**.

**Figure 1.**
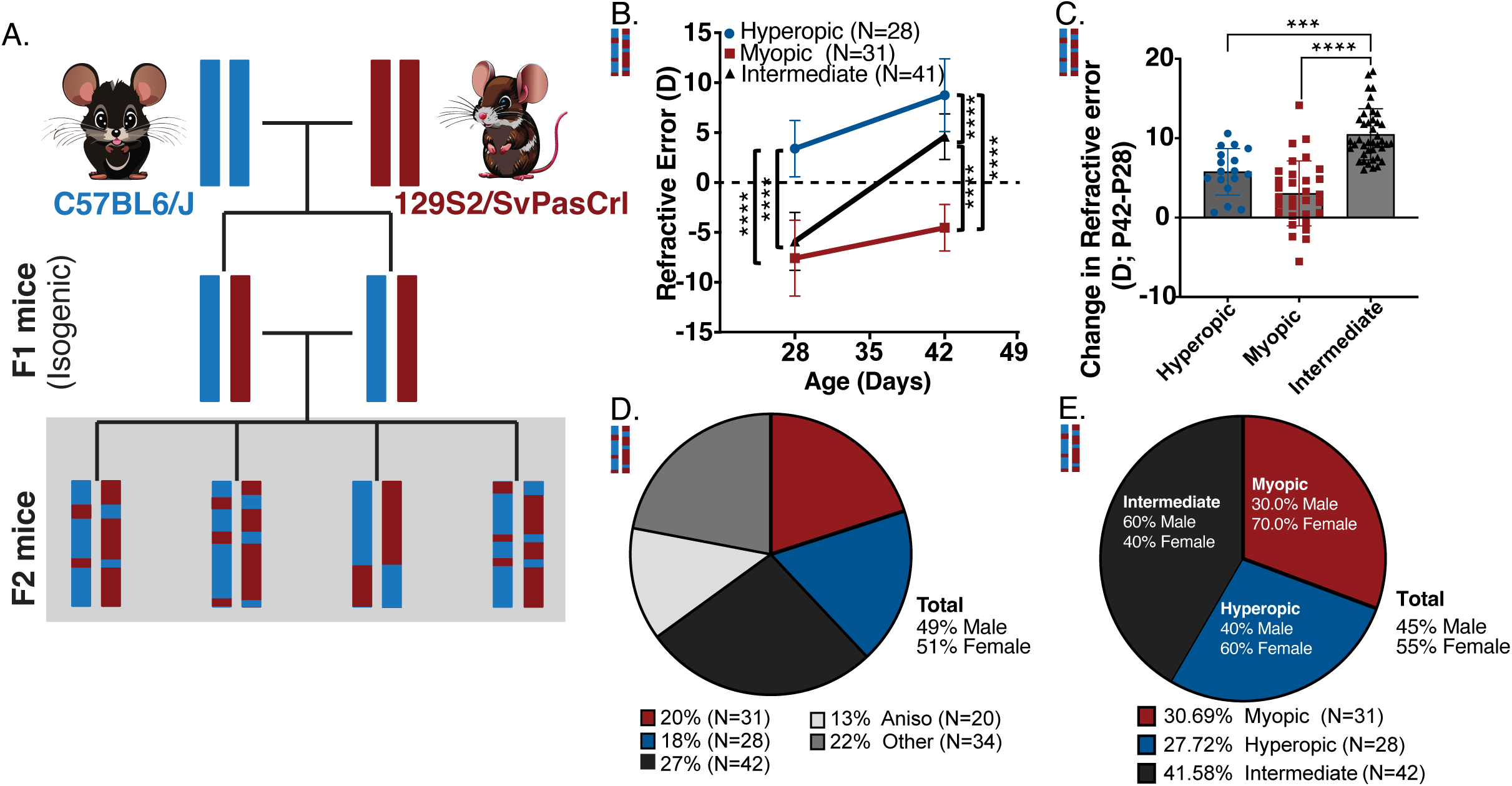
F2 outcrossed mice had hyperopic, myopic, and intermediate refractive development phenotypes. A) Schematic of the breeding strategy used to generate F2 mice. B) F2 mice sorted into distinct refractive error phenotypes of hyperopic (blue circle), myopic (red square), and intermediate (black triangle) F2 mice from P28 to P42. Two-way ANOVA with Sidak’s multiple comparisons test. C) The refractive error change from baseline (P42-P28) was largest in the intermediate versus myopic and hyperopic mice. Each data point represents one animal. Kruskal-Wallis test with multiple comparisons. D) The percentages of myopic, hyperopic, and intermediate mice are fairly evenly distributed, after excluding mice from the study due to anisometropia or other disqualifying characteristics. E) The proportion of male and female mice differed depending on the refractive phenotype. *p<0.05, **p<0.01,***p<0.001, and ****p<0.0001. mean±SD.

### Refractive error characteristics in F1 mice reveal key differences in inheritance

To further evaluate the inheritance patterns of refractive phenotypes, first-generation offspring (F1) were generated, and their refractive error was measured. F1 mice displayed only two phenotypes compared to the F2 generation of mice: myopic and intermediate (**Figure 2A**). At P28, myopic (-6.2±2.4D) and intermediate (-5.8±2.1D) mice had comparable refractive errors; however, the two groups diverged significantly by P42 (-1.0±0.8D vs +3.0±1.7D; 2-way ANOVA with multiple comparisons P=0.0005), with 81% of the mice falling into the intermediate phenotype and 19% into the myopic phenotype (**Figure 2B**). This phenotypic distribution ultimately indicates a dominant inheritance pattern for myopic refractive errors at earlier ages and a more complex genetic interaction that drives refractive eye growth with age.

**Figure 2.**
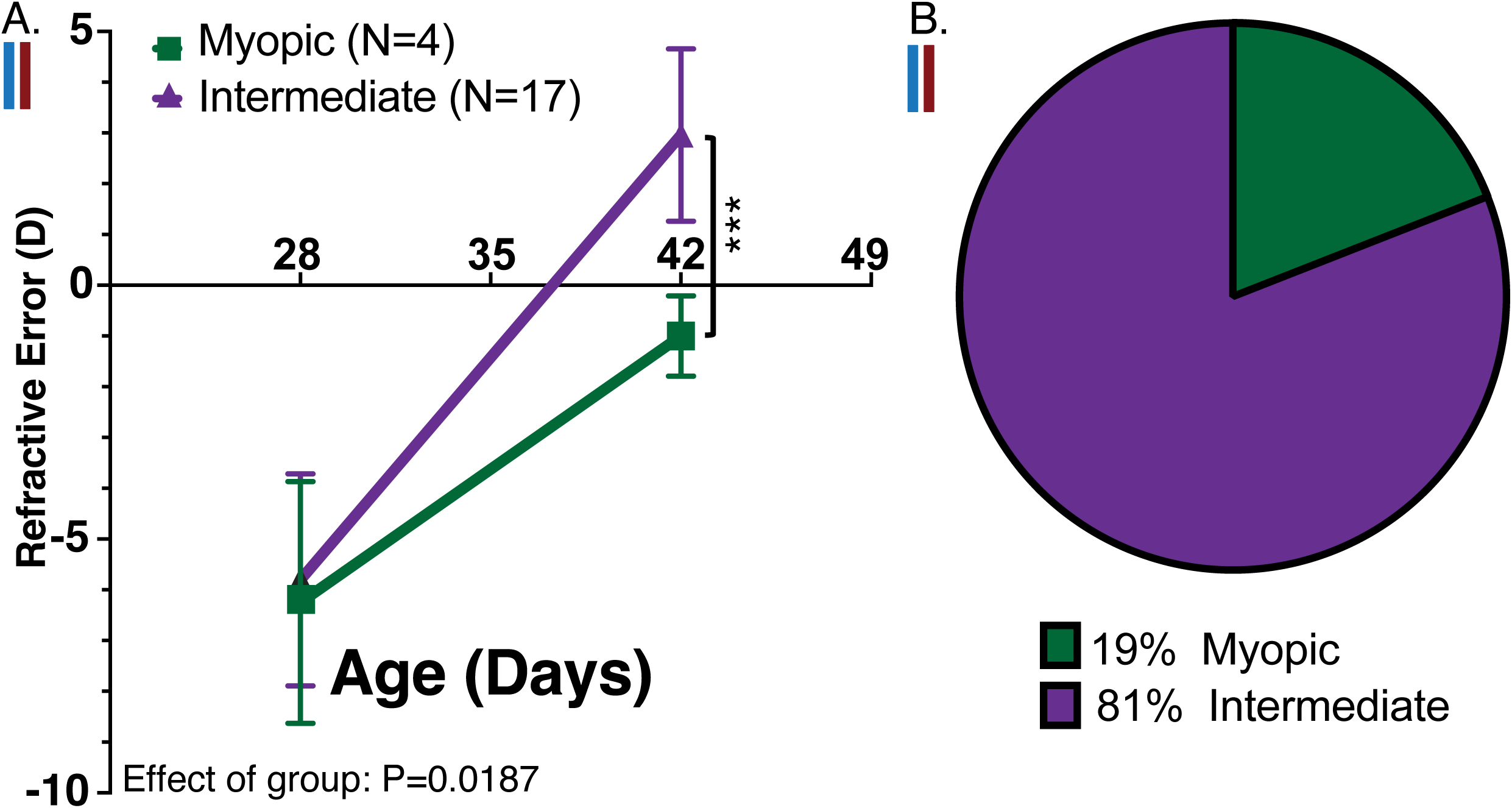
Myopic refractive error was the dominant inheritance pattern in F1 mice. A) F1 mice had similar myopic refractions that at P28 that then developed into two distinct groups at P42: myopic (green square) and intermediate (purple triangle) Two-way ANOVA with uncorrected Fisher’s least significant difference test. B) The majority of the F1 mice had the intermediate phenotype.. *p<0.05, **p<0.01,***p<0.001, and ****p<0.0001. mean±SD.

### Differences in axial growth rates and lens thickness contribute to refractive error differences between phenotypes

To determine whether axial parameters contribute to differences in refractive error between groups, we used SD-OCT to quantify them over time. While there were no significant effects of phenotype on axial length (**Figure 3A**), myopic mice showed significantly greater change in axial length between P28 and P42 than intermediate mice (**Figure 3B**; one-way ANOVA, P=0.002; Tukey’s multiple comparisons; P=0.0016). Similarly, there were no significant effects of phenotype on vitreous chamber depth at P28 or P42 (**Figure 3C**); however, intermediate mice exhibited a larger reduction in vitreous chamber depth from P28 to P42 compared to myopic (**Figure 3D**; one-way ANOVA P=0.01; Tukey’s multiple comparisons; P=0.04) and hyperopic mice (Tukey’s multiple comparisons; P=0.03). Interestingly, lens thickness of myopic mice was significantly thinner at P28 than hyperopic (**Figure 3E**; Mixed-effects analysis, P=0.04; multiple comparisons; P=0.02) and intermediate mice (multiple comparisons; P=0.03). Lens thickness was not different between phenotypes at P42, nor did the change from baseline significantly differ between groups (**Figure 3F**; Mixed-effects analysis; p=0.41). Refractive phenotype had no significant effect on any other axial parameters measured by SD-OCT (**Supplemental Figure 2A-J**). To evaluate the potential contributions of corneal curvature, keratometry was performed, but it revealed no significant differences (**Supplemental figures 2K, L).**

**Figure 3.**
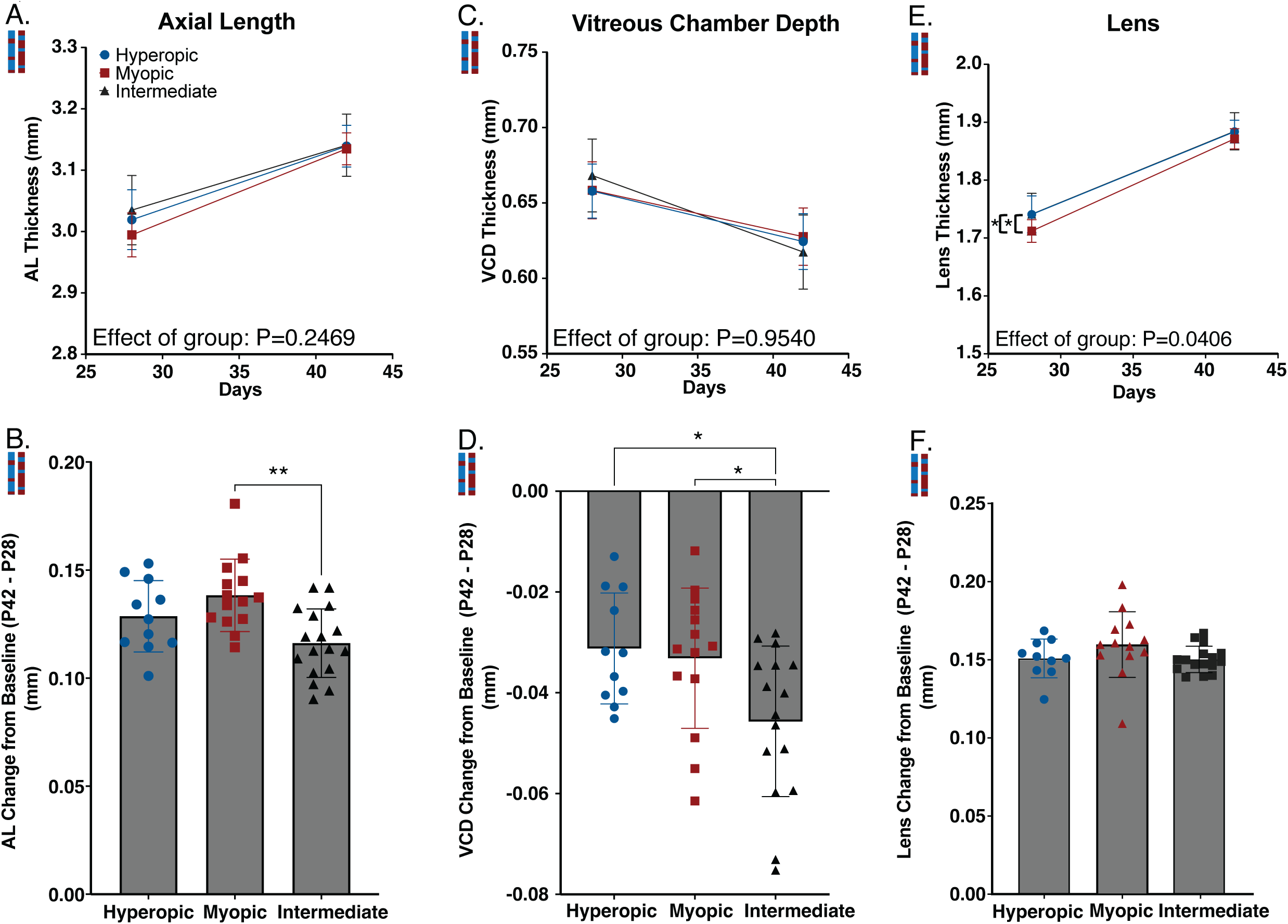
Differences in axial growth rates and lens thickness contributed to differences in refractive error between NORE groups. SD-OCT measurements of A) axial length (AL) at P28 and P42, B) change in AL from P28-P42, C) vitreous chamber depth (VCD) at P28 and P42, D) change in VCD from P28-P42, E) lens thickness (LT) at P28 and P42, F) change in LT from P28-P42. AL and VCD were significantly different in the intermediate phenotypes, suggesting a slowing of axial growth with age to become more hyperopia. A,C,E) Mixed-effects analysis. B,D,F) One-way ANOVA with Tukey’s multiple comparisons test, each data point represents one animal. ∗ P≤0.05, ∗∗ P≤0.01, mean±SD.

### Refractive phenotypes have similar retinal and visual function

No differences in photoreceptor function were observed, as quantified by a-wave amplitude (**Figure 4A**; mixed effects analysis; effect of phenotype P=0.11) or a-wave implicit time (**Figure 4B**; mixed effects analysis; effect of phenotype P=0.33). Furthermore, retinal ON pathway function was assessed using b-wave amplitude, as deficits in this pathway have been implicated in myopiaa^46–52^. The refractive phenotypes did not show significant differences in b-wave amplitude (**Figure 4C**; mixed-effects analysis; effect of phenotype P=0.45) or implicit time (**Figure 4D**; mixed-effects analysis; effect of phenotype P=0.76). Visual function was assessed by optomotor response to test spatial frequency threshold (**Figure 5A**) and contrast sensitivity (**Figure 5B**). Spatial frequency thresholds and contrast sensitivity did not differ significantly across groups. Collectively, these data indicate that retinal function is preserved in all refractive groups.

**Figure 4.**
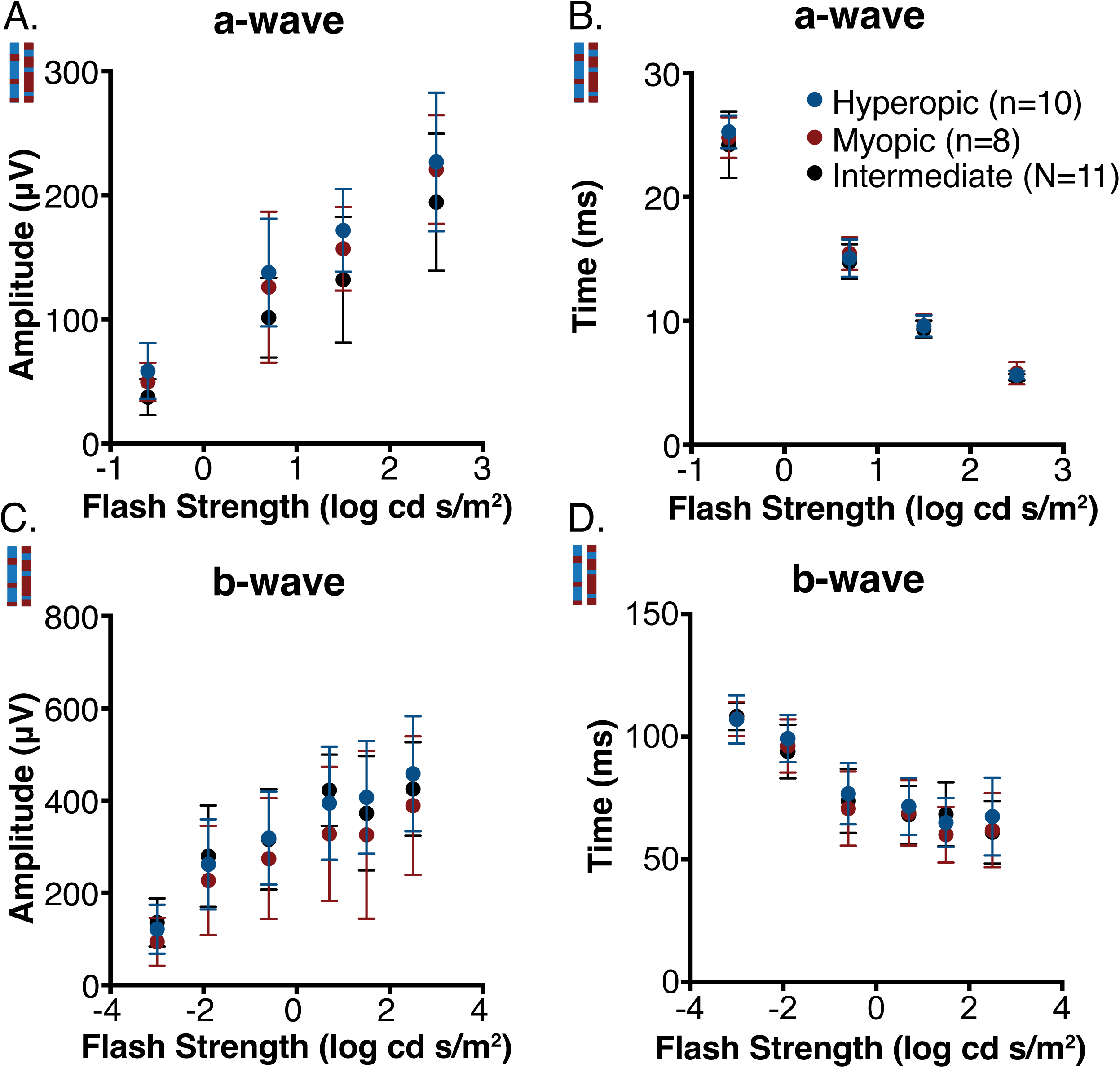
Retinal function was similar between NORE phenotypes. Quantification of scotopic ERG A) a-wave amplitude, B) a-wave implicit time, C) b-wave amplitude, and D) b-wave implicit time showed no differences. Mixed-effects analysis, mean±SD.

**Figure 5.**
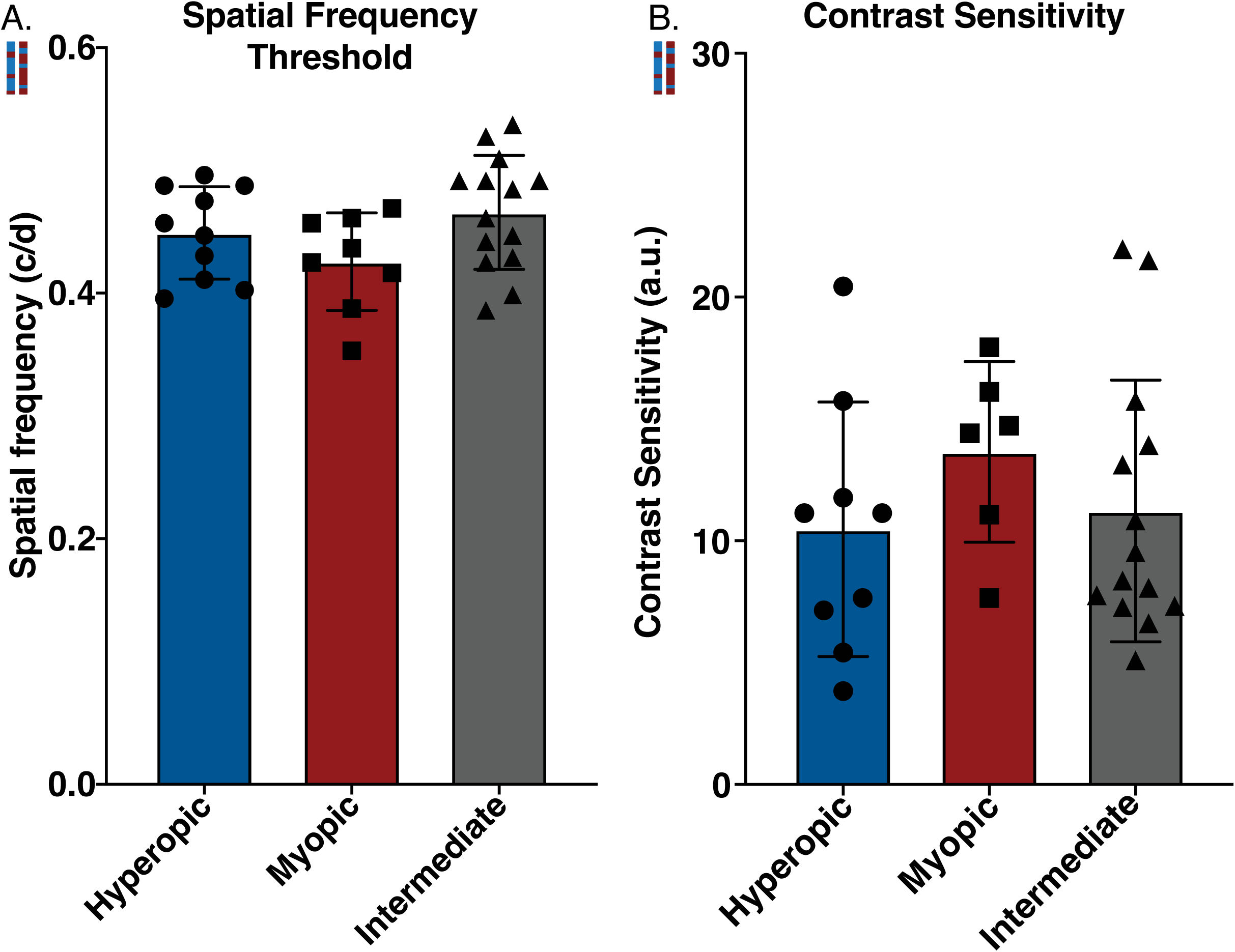
Spatial frequency thresholds and contrast sensitivity are similar between NORE phenotypes. Assessment of A) spatial frequency thresholds and B) contrast sensitivity by optomotor response (OMR) showed no significant differences between refractive phenotypes.

## Discussion

The NORE mouse model provides a powerful model of three naturally occurring refractive error phenotypes: hyperopic, myopic, and intermediate. It not only enables direct comparison between myopic and hyperopic phenotypes but also provides an intermediate group that displays a greater change in axial growth with age. While there is potential to directly compare different inbred strains, such as 129SvPasCrl and C57BL/6J, these comparisons are confounded by strain-specific genetic backgrounds. In contrast, the NORE model allows for the study of genetically unique mice while retaining the advantages of inbred strains, specifically, access to complete genome sequences and the ability to trace genomic loci to their originating strains. Ongoing and future work will leverage this feature to perform quantitative trait locus (QTL) analyses to identify genomic regions associated with phenotypes such as refractive error and assess the polygenic nature of myopia. This will provide further opportunity to perform targeted analysis of genetic contributions to refractive development.

A key advantage of the NORE model is the ability to compare phenotypes within the same litter of mice, providing critical internal environmental controls. It also provides the unique opportunity to probe different refractive phenotypes without requiring any intervention. Mutant mouse models have been instrumental in advancing our understanding of refractive development. For example, *Cx36* (*Gjd2*) mutant mice have provided insight into the gene most strongly associated with myopia in humans, according to GWAS studies^53–56^. Further*, nob* (no b-wave) alleles have provided critical information on the contributions of retinal ON signaling in myopia^49^ ^46,57–59^. However, their association with syndromic myopia complicates their interpretation in the context of normal refractive development patterns. In contrast, the NORE model better mimics the multifactorial nature of myopia in humans, in which refractive errors arise from complex gene network -environment interactions rather than single-gene mutations. An additional strength of the NORE model is the ability to group or categorize the F2 population in different ways to investigate traits of interest. For example, rather than categorizing mice as hyperopic, myopic, or intermediate, as we have done here, analysis could be performed on mice based on the rate of change in RE or the rate of axial elongation. In genomic studies, these mice can be evaluated separately to identify unique identifiers of the phenotype of choice. The timeframe evaluated in NORE mice, approximately P28 to P42, is consistent with many studies conducted in mice using LIM or FD models^27,46,50,60^. Thus, these results provide an opportunity to directly compare the results from NORE mice to mice with induced experimental refractive errors.

Phenotyping performed in this study suggests that the shift from negative to positive refractive error in the intermediate group may be partially explained by slower axial growth with age compared with myopic and hyperopic mice. Surprisingly, myopic mice had thinner lenses than hyperopic and intermediate mice at P28, which likely contributes to the respective differences in refractive error. Even though we did not observe significant alterations in scleral thickness on OCT, these changes in axial elongation rates warrant further investigation of potentially underlying scleral properties and retinoscleral signaling events. Previous work in the mouse has shown measurable differences in matrix composition and biomechanics despite negligible differences in thickness^61,62^. Analysis of retinal and visual function via ERG and OMR revealed no deficits in retinal or visual function between NORE refractive phenotypes. This is advantageous as it provides an opportunity to evaluate the refractive phenotypes without confounds of visual or retinal dysfunction to the development of refractive errors.

A further consideration with NORE is the different prevalences of myopic and hyperopic phenotypes in male and female mice. This presents an opportunity to investigate previously reported differences in the prevalence of myopia between males and females in clinical studies, without confounding social factors^63^. Moving forward, our genetic studies will consider this factor to try to identify what may be driving this effect. It is important to note that this should be considered when using this model by including both sexes in experiments.

In conclusion, the NORE mice provide a novel model for the myopia field which can be used in future studies to determine genetic links to refractive phenotypes. Additionally, NORE mice can be exposed to various visual conditions to elucidate the complex gene-environment interactions that govern refractive eye growth, thereby identifying mechanisms of myopia that can inform the development of therapeutic options to prevent myopia.

## Acknowledgements

We would like to thank Micah Chrenek, the ophthalmology research lab manager. The Georgia Clinical & Translational Science Alliance of the National Institutes of Health provided additional support under Award Number UL1TR002378.

## Figure legends

**Supplemental Figure 1.**
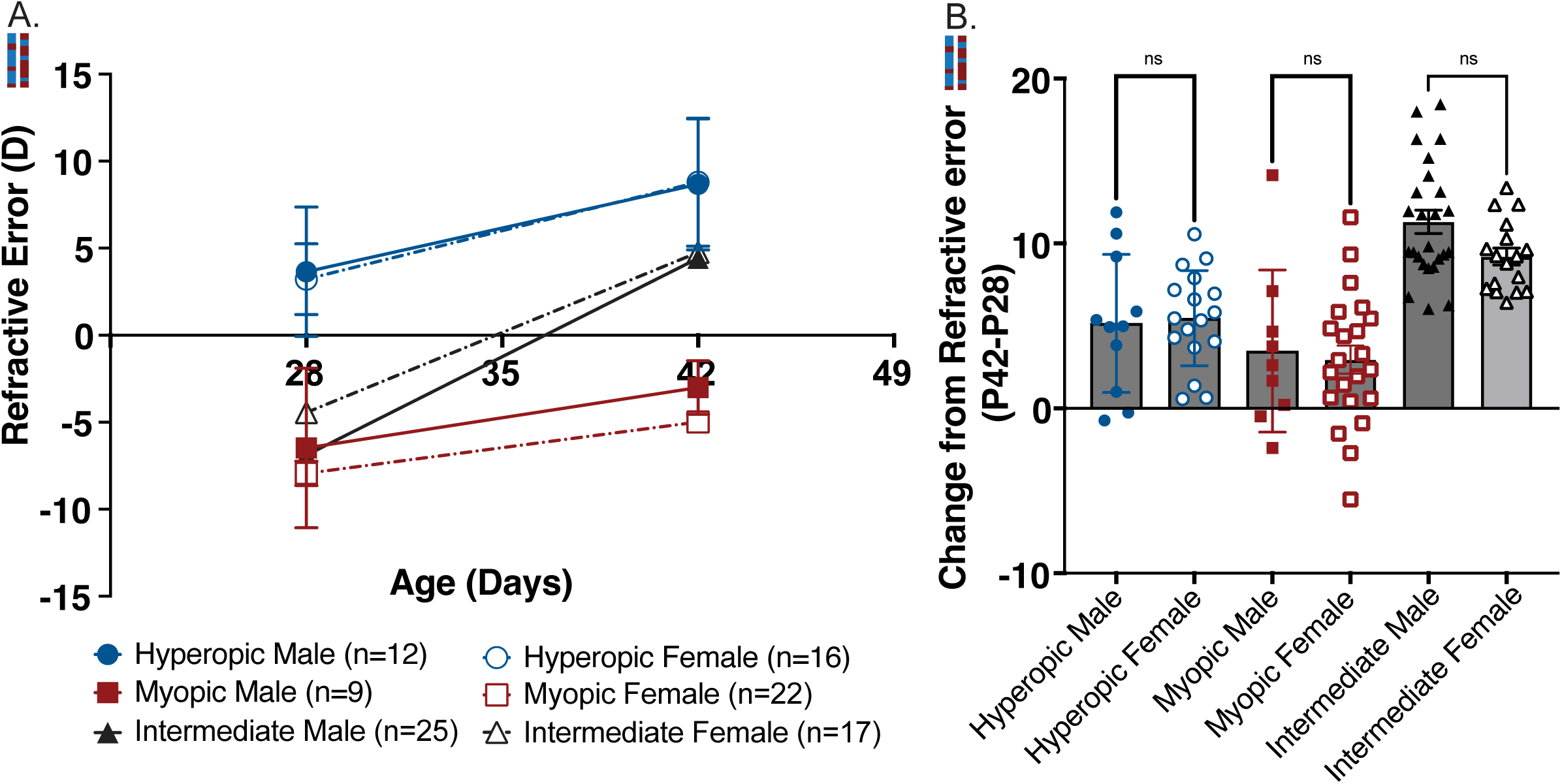
NORE mice refractive phenotypes were not dependent on sex.. A) Refractive errors of hyperopic, myopic, and intermediate animals showed no differences between male (filled symbols) and female (open symbols) mice. B) The change in refractive error from baseline (P42-P28) in hyperopic, myopic, and intermediate animals was similar between male (filled symbols) and female (open symbols) mice. A) Two-way ANOVA with multiple comparisons (Effect of sex between myopic males and females, p=0.0586). B) Brown-Forsythe and Welch ANOVA with multiple comparisons, each data point represents one animal. mean±SD.

**Supplemental Figure 2.**
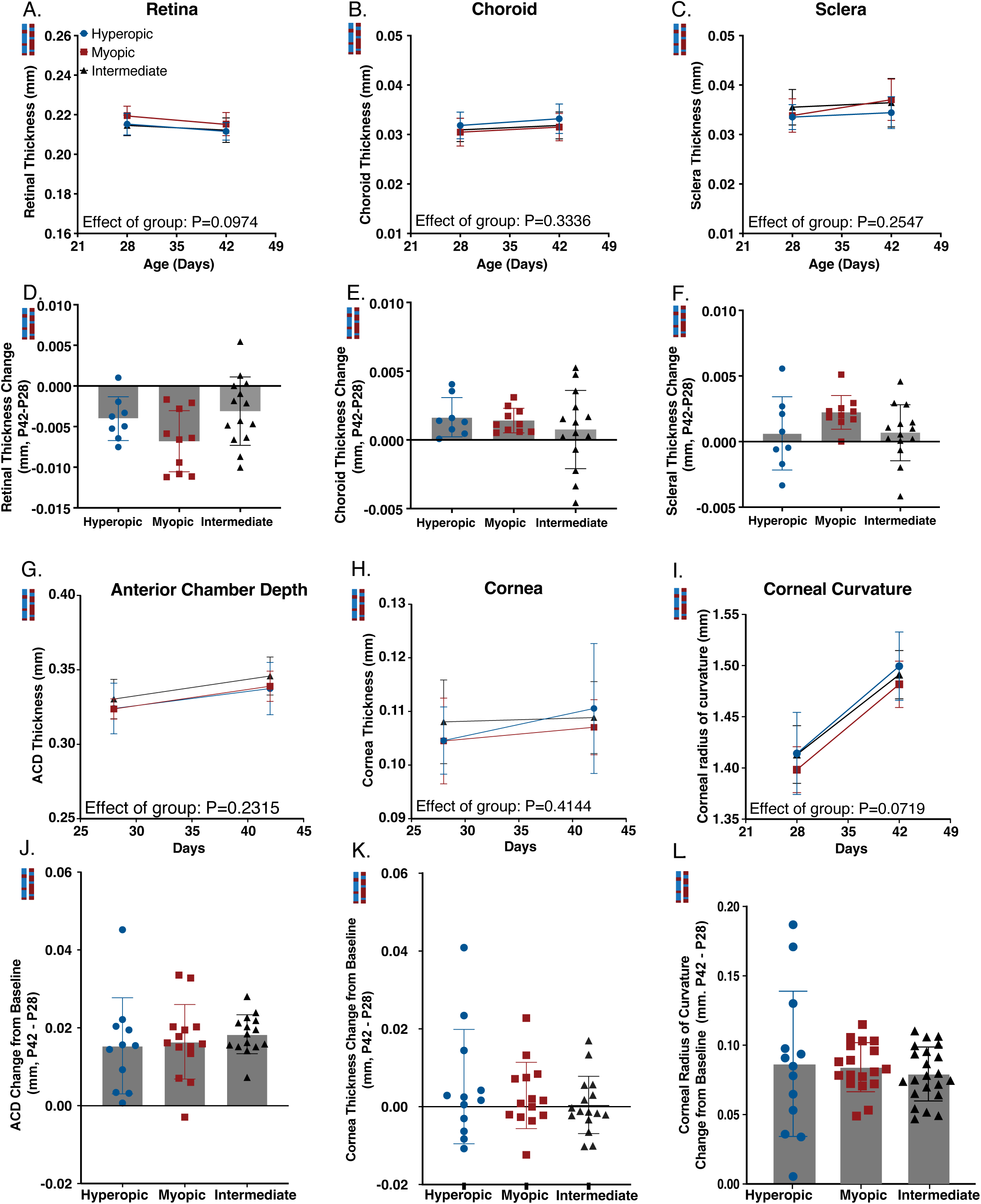
Additional biometry measurements from NORE mice revealed no differences in ocular anatomy. SD-OCT measurements across age (A-B, G-I) and change from baseline (D-F, J-L) showed similar results for all refractive phenotypes for A,D) retinal thickness (RT), B,E) choroidal thickness (ChT) at P28 and P42, C,D) scleral thickness (ST), G, J) anterior chamber depth (ACD), H,K) corneal thickness (CT) and I, L) Corneal radius of curvature (CC) Mixed-effects analysis. B,D,F,H,J,L) one-way ANOVA with Tukey’s multiple comparisons test, each data point represents one animal. ∗ P≤0.05, ∗∗ P≤0.01, ∗∗∗ P≤0.001 mean±SD.

